# Combinatorial phosphorylation modulates the structure and function of the G protein gamma subunit in yeast

**DOI:** 10.1101/2020.06.09.143123

**Authors:** Zahra Nassiri Toosi, Xinya Su, Shilpa Choudhury, Wei Li, Yui Tik Pang, James C. Gumbart, Matthew P. Torres

**Author notes:** To Whom Correspondence Should be Addressed: Engineered Biosciences Building, School of Biological Sciences, Georgia Institute of Technology.

## Abstract

Protein intrinsically disordered regions (IDRs) are often targets of combinatorial post-translational modifications (PTMs) that serve to regulate protein structure and/or function. Emerging evidence suggests that the N-terminal tails of G protein γ subunits – essential components of heterotrimeric G protein complexes – are intrinsically disordered, highly phosphorylated governors of G protein signaling. Here, we demonstrate that the yeast Gγ Ste18 undergoes combinatorial, multi-site phosphorylation within its N-terminal IDR. Phosphorylation at S7 is responsive to GPCR activation and osmotic stress while phosphorylation at S3 is responsive to glucose stress and is a quantitative indicator of intracellular pH. Each site is phosphorylated by a distinct set of kinases and both are also interactive, such that phosphomimicry at one site affects phosphorylation on the other. Lastly, we show that phosphorylation produces subtle yet clear changes in IDR structure and that different combinations of phosphorylation modulate the activation rate and amplitude of the scaffolded MAPK Fus3. These data place Gγ subunits among the growing list of intrinsically disordered proteins that exploit combinatorial post-translational modification to govern signaling pathway output.

## Introduction

Combinatorial post-translational modification (PTM) of intrinsically disordered protein termini has emerged as a prominent regulatory feature in protein biology. Indeed, several well-characterized cases are known to exist, in which the core regulatory element is defined by three criteria: the N- or C-terminal location of an intrinsically disordered region (IDR), the dynamic and combinatorial modification of amino acid sidechains within the IDR, and the propensity for such changes to alter protein interactions and functional output. Potent examples include the N-terminal tails of histone proteins, which undergo complex combinatorial PTM that governs eukaryotic gene transcription (*1*); the C-terminal domain of RNA polymerase II, which undergoes dynamic site-specific phosphorylation that governs transcription initiation and elongation (*2*); the C-terminal tails of tubulin, which undergo combinatorial modification that coordinates microtubule-end protein interactions (*3*); the C-terminal tails of G protein-coupled receptors (GPCRs), which undergo bar-code-like phosphorylation that coordinates protein interactions that govern pathway selectivity in G protein signaling (*4–6*); and other well-known examples that are also conserved across eukaryotes from yeast to humans (*7, 8*). In this work, we tested the hypothesis that heterotrimeric G protein gamma subunits (Gγ), small yet essential GPCR signal transducing proteins with N-terminal IDR phosphorylation hotspots, undergo regulatory combinatorial phosphorylation similar to that of classically defined PTM-regulated terminal IDRs.

Heterotrimeric G proteins (G proteins), an evolutionarily conserved group of protein families consisting of Gα, Gβγ, and Gγ subunits, function as transducers of extracellular signals such as light, hormones, and neurotransmitters that can activate 7-transmembrane G protein-coupled receptors (GPCRs) embedded within the plasma membrane of eukaryotic cells (*9*). Activation of the receptor on its extracellular surface stimulates a conformational change in the Gα subunit that promotes guanine nucleotide exchange and facilitates its dissociation from Gβγ – an obligate heterodimer of Gβ and Gγ subunits. Once dissociated, both Gα and Gβγ are free to interact with and modulate the activity of downstream protein effectors that control the production of second messenger signals, the activation of kinases, and the coordination of macro-molecular events that constitute the cellular response to the stimulus. Signaling continues until Gα hydrolyzes GTP back to GDP and re-enters a conformation that sequesters Gβγ – a process that is accelerated by their interaction with regulators of G protein (RGS) proteins (*10*).

As the smaller member of the obligate heterodimeric Gβγ complex, Gγ subunits are often thought to have limited functionality as membrane anchors for Gβ subunits – a function mediated through lipidation of C-terminal cysteine residues and through coiled-coil interactions with Gβ N-terminal residues (*11–13*). However, recent evidence suggests that the N-terminal regions of Gβ and Gγ, which are proximal to each other in the Gβγ complex, play an important role in Gβγ-dependent effector signaling. In the budding yeast *Saccharomyces cerevisiae* (yeast), the effector binding sites on Gβγ are located in N-terminal residues in Gβ_Ste4_ (*14, 15*), which overlap with the same region in human Gβ_1_γ_2_ that inhibit phospholipase C β2 activity in mammals (*16*). Mutation near this region has also been found to modulate mammalian Gβγ-dependent activity of adenylyl cyclase (AC) 5 and 6 *in vitro* (*17*).

The extreme N-termini of Gγ subunits are intrinsically disordered phosphorylation hotspots shown to govern Gβγ/effector interactions and activity (*18, 19*). Meta-proteomic informatics analysis has revealed that Gγ subunits harbor 2-fold greater modification density (average number of PTMs per total protein length in residue number) than Gα or Gβ subunits, RGS proteins, or GPCRs (*19, 20*). Moreover, all Gγ subunits throughout Eukarya harbor an N-terminal IDR. Within this IDR, which averages between 7-13 residues in length, most Gγ proteins harbor at least two phospho-acceptor residues and approximately 60% of all experimentally observed Gγ phosphorylation – 12 times more than what is observed outside the N-terminal IDR (*19*). Previous work in mammals first demonstrated that protein kinase C (PKC)-dependent phosphorylation within this hotspot in Gγ_12_ inhibits AC_2_ stimulation by Gβ_1_γ_12_ *in vitro* (*21*), and is required for normal fibroblast motility *in vivo* (*22*).

N-terminal Gγ phosphorylation is also required for G protein signaling in yeast. Budding yeast Gγ, Ste18, undergoes MAPK-dependent feedback phosphorylation within its N-terminal IDR in response to pheromone-dependent activation of the pheromone GPCR, Ste2 (*18*). The Ste18 N-terminal IDR harbors three phospho-acceptor residues (T2, S3, and S7) and nullification of all three sites by mutation to alanine (S/T-A), in combination with hyperactivating mutations in the Gβγ-effector and MAPK scaffold, Ste5, results in rapid and prolonged bulk-accumulation of the scaffold at the plasma membrane and subsequent ultra-fast and elevated activation of the terminal MAPK, Fus3. In contrast, phosphomimic mutation of the same residues to glutamate (S/T-E) results in wild-type Ste5 recruitment and MAPK activation. These data suggested to us that, like other PTM-regulatory terminal IDRs, phosphorylation within the N-termini of Gγ subunits can serve to govern their protein interactions and functional output.

Here we investigated the hypothesis that Gγ subunit N-termini undergo combinatorial regulatory phosphorylation. Using the canonical yeast Gγ subunit, Ste18, as a model, we measured the sensitivity of N-terminal phospho-acceptor sites T2, S3, and S7 to a variety of stimuli including GPCR activation by yeast mating pheromone, osmotic stress, glucose stress, and acid stress. Of these three sites, we find that S7 is phosphorylated in response to pheromone-induced GPCR activation and osmotic stress whereas S3 is phosphorylated in a quantitative manner that is indicative of intracellular pH. We find that phosphorylation at S3 and S7 is also interdependent, such that nullifying or mimicking mutations at either site affects phosphorylation at the other. Lastly, we show that different phosphorylation states have subtle yet clear differences in IDR structure *in vitro* as well as distinct functional effects on MAPK activation *in vivo*. Together, this new evidence places G protein γ subunits in the group of proteins governed by combinatorial PTM-regulated IDRs.

## Results

### The N-terminal IDR of Ste18 undergoes stimulus- and site-specific phosphorylation

The N-terminal IDR of Ste18, Ste18^Nt^, has been shown to undergo regulatory phosphorylation in response to pheromone-dependent GPCR activation (*18*), but was originally first discovered by largescale phospho-proteomics analysis of yeast undergoing osmotic stress (*23*). Considering its regulatory role in the pheromone response pathway, we hypothesized that the tail was a hotspot for phosphorylation-dependent pathway crosstalk. To test this hypothesis, we utilized a phosphorylation-dependent electrophoretic mobility shift assay optimized for Ste18 to measure phosphorylation in response to common cellular stressors known to crosstalk with the pheromone pathway – namely osmotic stress, nutrient/glucose stress, and pH stress (*24*). Previously, pheromone-dependent phosphorylation was shown to correspond to the upper (slower migrating form) of two Ste18-specific immunoblot bands that was sensitive to phosphatase treatment (*18*). We first confirmed this result, showing that Ste18 phosphorylation occurs rapidly in the presence of α-factor mating pheromone and persists throughout the exposure period of 90 minutes (Figure 1A). Next, we exposed cells to osmotic shock with 0.75M potassium chloride, which has been shown to temporarily activate the osmotic stress pathway within 15 minutes (*25*). Ste18 phosphorylation was again clearly evident although distinctly different from the pheromone response, occurring to a lesser extent and for a much shorter period of time ranging from 15 to 45 minutes and then sharply decreasing (Figure 1B). In both pheromone and osmotic stress conditions we also noticed a variable degree of basal phosphorylation in untreated cells (t=0), suggesting that there is a low level of constitutive phosphorylation that occurs in the absence of stimulus or stress (Figure 1A,B).

**Figure 1.**
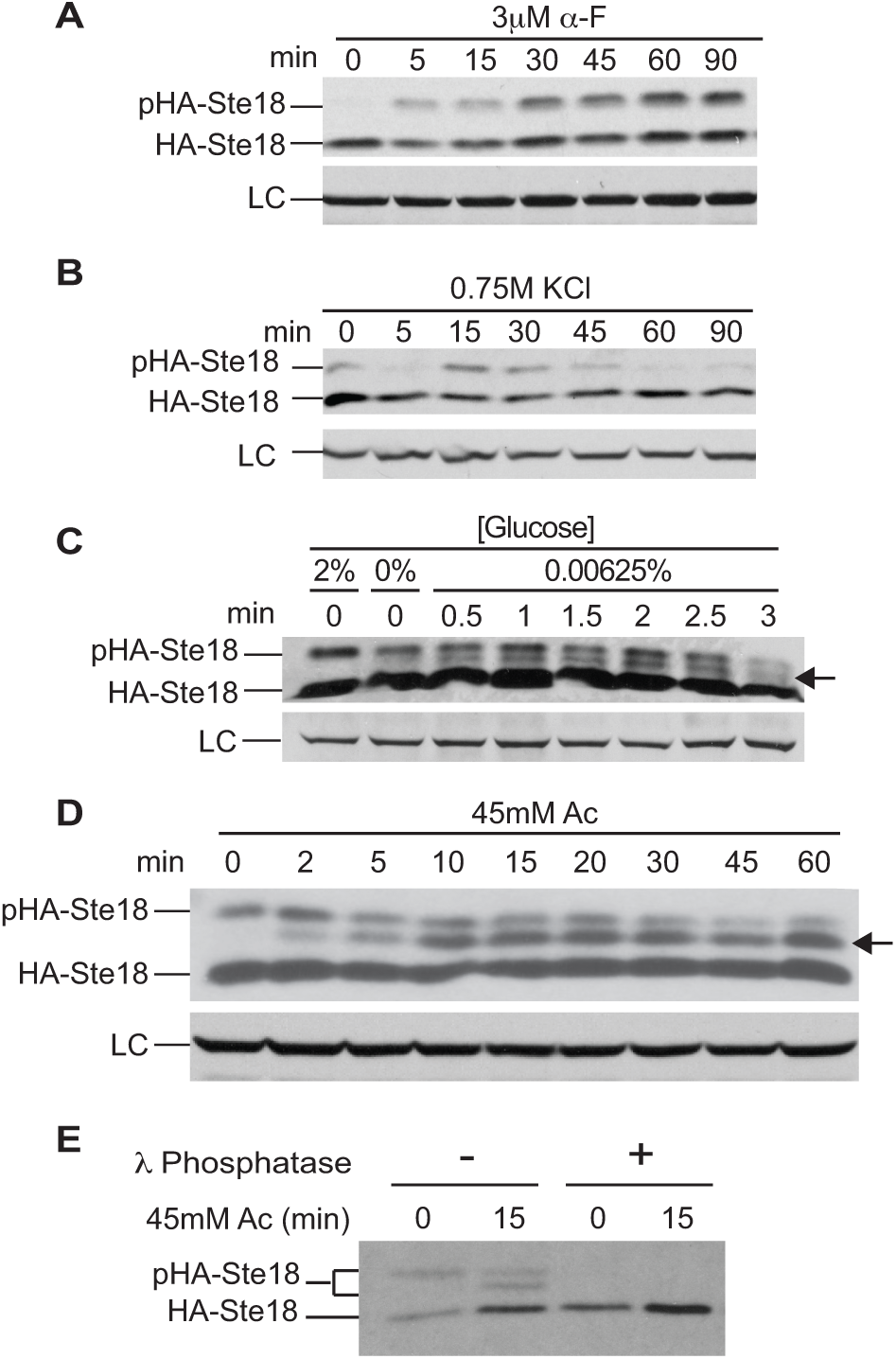
Ste18 is phosphorylated in response to cell stress. Phosphorylation dependent electrophoretic mobility of HA-Ste18 was monitored by immunoblotting with anti-HA antibody and in response to: (A) 3μM α-factor mating pheromone; (B) 0.75M potassium chloride; (C) 0%, 0.00625% (low) and 2% (standard) glucose; and (D) 45mM acetic acid (Ac). (E) Protein extracts of cells exposed to acid stress (45mM acetic acid for 15 minutes) were treated with or without alkaline phosphatase (AP) to establish the dependence of each mobility-shifted band on phosphorylation. (LC) loading control corresponding to yeast GAPDH; (pHA-Ste18) phosphorylated HA-Ste18.

Unlike pheromone stimulation or osmotic stress, we observed a unique banding pattern we had not seen previously in cells rapidly shifted from normal 2% glucose to low glucose media (0.00625%), whereby a unique band (middle band) between the phosphorylated and non-phosphorylated protein appeared shortly after the onset of the stress (0.5 min) representing ∼2-10% of total Ste18 levels (Figure 1C). This pattern was only observed under low glucose shock, and did not occur when cells were transferred to media completely lacking glucose, consistent with a previous report that low glucose but not the absence of glucose stimulates phosphorylation of some G proteins (*26*). Considering that glucose deprivation results in acidification of the intracellular environment and it is this factor that drives phosphorylation of G protein α subunits in yeast (*27*), we tested whether the 3-band pattern in Ste18 immunoblots could be induced by exposure of cells to acetic acid and whether this pattern was the result of phosphorylation. Indeed, we found that treatment of cells with 45mM acetic acid resulted in the same mobility shifted bands observed under glucose stress (Figure 1D), and that both the upper and middle bands corresponded to phosphorylated forms of the protein that were sensitive to phosphatase treatment (Figure 1E). Taken together, these data indicate that Ste18 phosphorylation is activated in response to a wide range of stimuli including both GPCR-dependent and independent signals; and moreover, that phosphorylation occurs at more than one site.

### Phosphorylation of Ste18^Nt^ occurs on two distinct phosphosites – S3 and S7

Based on prior mass spec evidence (*23*), we hypothesized that phosphorylation would occur on two sites within Ste18^Nt^, which contains three potential phospho-acceptor residues: T2, S3, and S7 (Figure 2A). To test this hypothesis, we substituted each acceptor residue in turn with either alanine (phospho-null) or glutamic acid (phospho-mimic) through point mutations introduced into the yeast genome. The resulting mutant yeast strains were exposed to either acetic acid or α-factor mating pheromone and endogenous levels of Ste18 proteins were evaluated by electrophoretic mobility analysis. Substituting T2 with either alanine (T2A) or glutamic acid (T2E) had no effect on the electrophoretic mobility of Ste18 in either condition, suggesting it was not a site of phosphorylation under these conditions (Figure 2B,C). Under acid stress, in which both middle and upper bands can be simultaneously detected, alanine substitution of S3 (S3A) resulted in a complete loss of the middle band while alanine substitution of S7 (S7A) resulted in a complete loss of the upper band (Figure 2B). Furthermore, nullifying phosphorylation at both S3 and S7 eliminated all phosphorylation-dependent mobility shifts (Figure S1). These results confirmed the identity of the middle band as phospho-S3 (pS3) and the upper band as phospho-S7 (pS7). These assignments were further confirmed by glutamate substitution of either site (S3E or S7E), which slowed the mobility of the non-phosphorylated lower band (np) in a manner reflective of the relative differences in electrophoretic mobility of pS3 and pS7 bands, respectively (Figure 2B). Consistent with these assignments, we found that the upper band (corresponding to pS7) increases in response to pheromone stimulation, is completely lost for cells expressing Ste18^S7A^, and is mimicked by the S7E mutation (Figure 2C). Taken together, these data show that the electrophoretic mobility of Ste18 reflects the phosphorylation state of two distinct residues in Ste18^Nt^: S3, a pH and glucose-sensitive phosphosite; and S7, a pheromone and osmotic stress-sensitive phosphosite.

**Figure 2.**
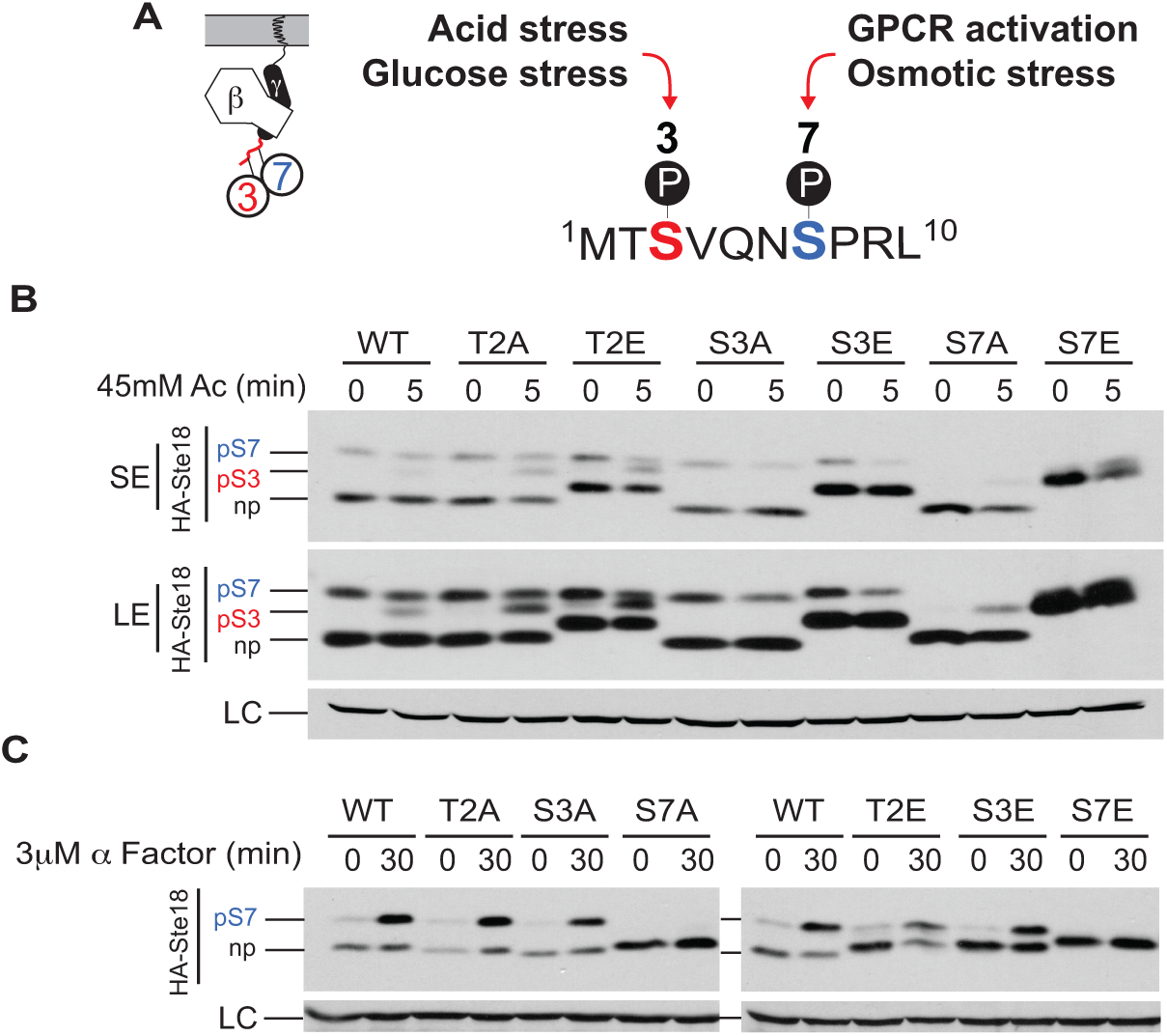
pH and pheromone dependent phosphorylation occur at unique phosphorylation sites in Ste18^Nt^. (A) Schematic summary of site-specific phosphorylation and corresponding stimuli across the intrinsically disordered N-terminal tail of Ste18. (B) Cells harboring single phosphosite mutations were treated with 45mM acetic acid (Ac) for 5 minutes or (C) with 3 μ M pheromone (α Factor) for 30 minutes followed by immunoblot analysis with anti-HA antibody. Phosphorylation on Ste18^Nt^ at Ser-7 (pS7) or Ser-3 (pS3), non-phosphorylated Ste18 (np). (LC) loading control corresponding to yeast GAPDH. (SE), short exposure; (LE), long exposure.

### Ste18^pS3^ and Ste18^pS7^ are inversely regulated in response to intracellular acidification

The characteristics of acid stress-dependent phosphorylation of Ste18^S3^ – discovered here for the first time – have never before been described. We hypothesized that changes in the phosphorylation of Ste18^Nt^ upon acid stress were the direct result of changes in intracellular pH (pH_i_), which should manifest as a kinetic and concentration-dependent correlation between pS3 and extracellular acid stress. To test this hypothesis, we monitored changes in the phosphorylation of Ste18^Nt^ relative to changes in pH_i_ across time and concentration of extracellular acetic acid stress. Intracellular pH was monitored using plasmid-borne expression of pHluorin – a pH-sensitive green fluorescent protein for which the ratio of fluorescence emission in response to excitation at 395nm and 480nm provides a direct readout of the pH environment inside the cytoplasm of cells (*28, 29*). Under non-buffered conditions, in which acetic acid is added directly to cell culture media without the addition of a buffer, we observed an instantaneous drop in pH_i_ from pH 7.2 to pH 4.5, which did not recover over the remainder of the 90-minute experiment (Figure 3A). Concomitantly, pS3 increased from 0 to ∼20% of total protein levels within ∼15 minutes (t_1/2_ < 5 min) and remained stable for the remaining duration of the experiment (Figure 3B,C). Interestingly, we observed that pS7 decreased with nearly identical kinetics over the same time period (identical t_1/2_), dropping from ∼30% to ∼5% of total Ste18 levels. Beyond 1 hour, no further changes in phosphorylation were observed, consistent with the highly stable pH_i_ observed in these cells.

**Figure 3.**
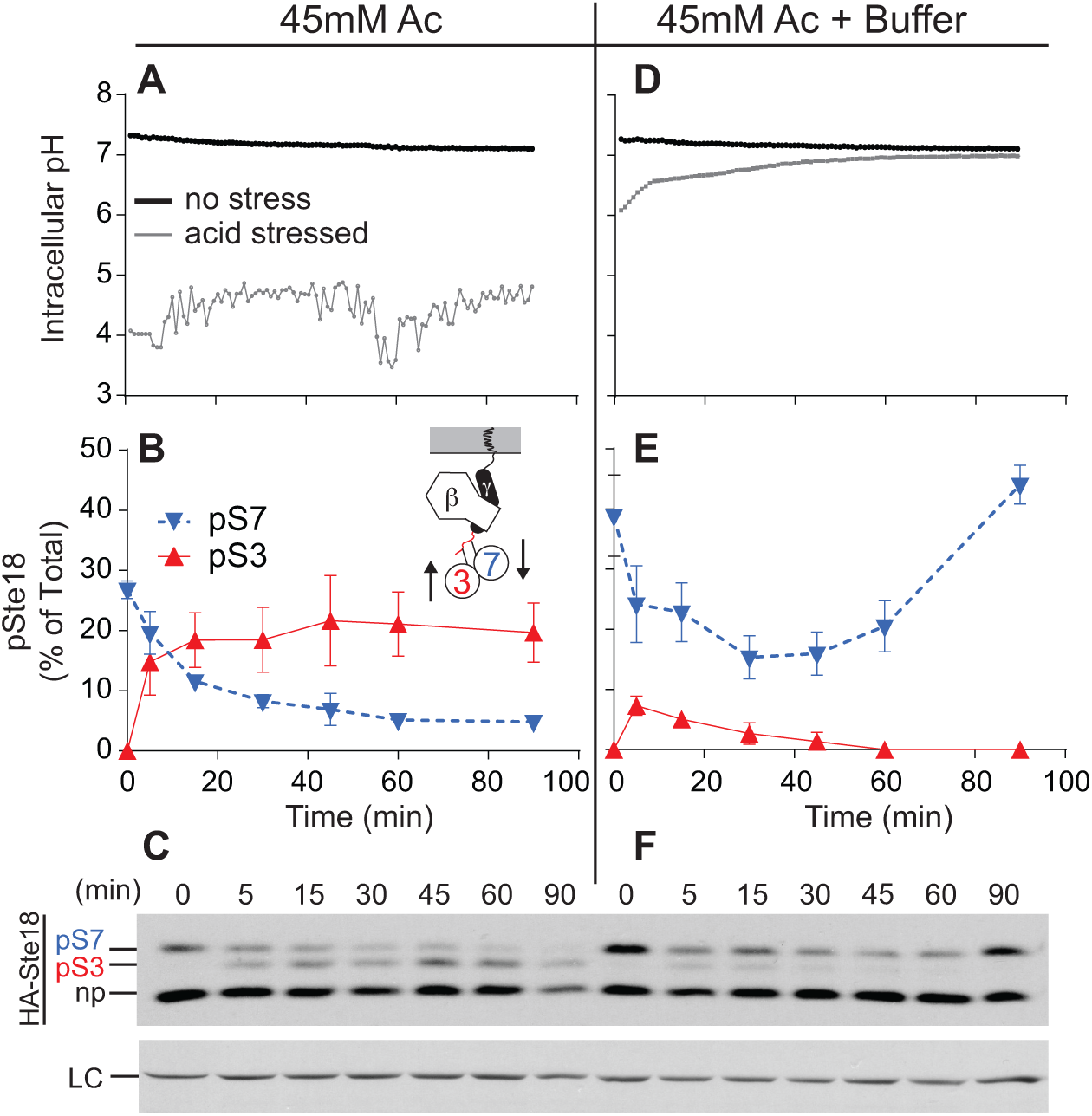
Ste18^pS3^ and Ste18^pS7^ are inversely regulated in response to intracellular acidification. Wild type *BY4741* yeast cells were transformed with a pH-sensitive pHluorin probe followed by timed exposure to 45mM acetic acid and analysis of intracellular pH (A) and site-specific Ste18 phosphorylation percentage (B,C). The same cell type was exposed to 45mM acetic acid in the presence of a buffer and the same measurements taken for comparison (D-F). Results are the mean ± SD (n=4). Ste18 phosphorylation at Ser-7 (pS7) or Ser-3 (pS3), and non-phosphorylated Ste18 (np). (LC) loading control corresponding to yeast GAPDH.

The addition of a buffer to yeast cell culture media has been shown previously to support the inherent ability of yeast cells to restore pH_i_ after encountering extracellular acid stress, mediated by the proton pump Pma1 (*27*). Exploiting this feature of yeast, we next asked whether under buffered conditions the phosphorylation state of Ste18 would be restored to its original levels after cells adapt to pH stress, which would indicate that the phosphorylation state is regulated by pH, rather than acetic acid specifically. Consistent with previous observations (*27*), addition of acetic acid to 45mM final concentration in the presence of a buffer had much less effect on pH_i_, resulting in an instantaneous drop from pH ∼7.2 to 6.0 followed by an hour-long adaptation period where pH_i_ was restored back to the level observed in control cells (Figure 3D).

Concomitant with the kinetics and amplitude of pH_i_ changes, phosphorylation of S3 was diminished when compared to the un-buffered experiment, increasing from 0 to only 7% within 5 minutes and then decreasing back to an undetectable level within 60 minutes (Figure 3E,F). As we had observed in un-buffered experiments the response of pS7 mirrored that of pS3, dropping only slightly from 40% to 20% by 30 minutes and then rebounding back to original levels by 90 minutes. These results demonstrate that Ste18^Nt^ undergoes dynamic multi-site phosphorylation coordinated in response to fluctuations in pH_i_.

Considering that Ste18^Nt^ phosphorylation was tightly coordinated with fluctuations in pH_i_ observed under buffered stress conditions, we asked whether it may be a quantitative indicator of the intracellular effects of acid stress. To test this hypothesis, we directly compared the fluorometric response of pHluorin with S3 phosphorylation across a concentration range of acetic acid doses added to yeast cell cultures without buffer. Indeed, we found that pS3 exhibited a sigmoidal dose response to increasing concentrations of acetic acid (Hill slope = 5.924), indicating that Ste18^Nt^ is cooperatively ultra-sensitive to pH_i_ (Figure 4). The pHluorin fluorescence response confirmed the decrease in pH_i_ with increasing acetic acid concentration but saturated beyond 40mM, above which only the phosphorylation percentage of pS3 was indicative of changing pH_i_. We conclude that Ste18^pS3^ is a quantitative indicator of intracellular pH.

**Figure 4.**
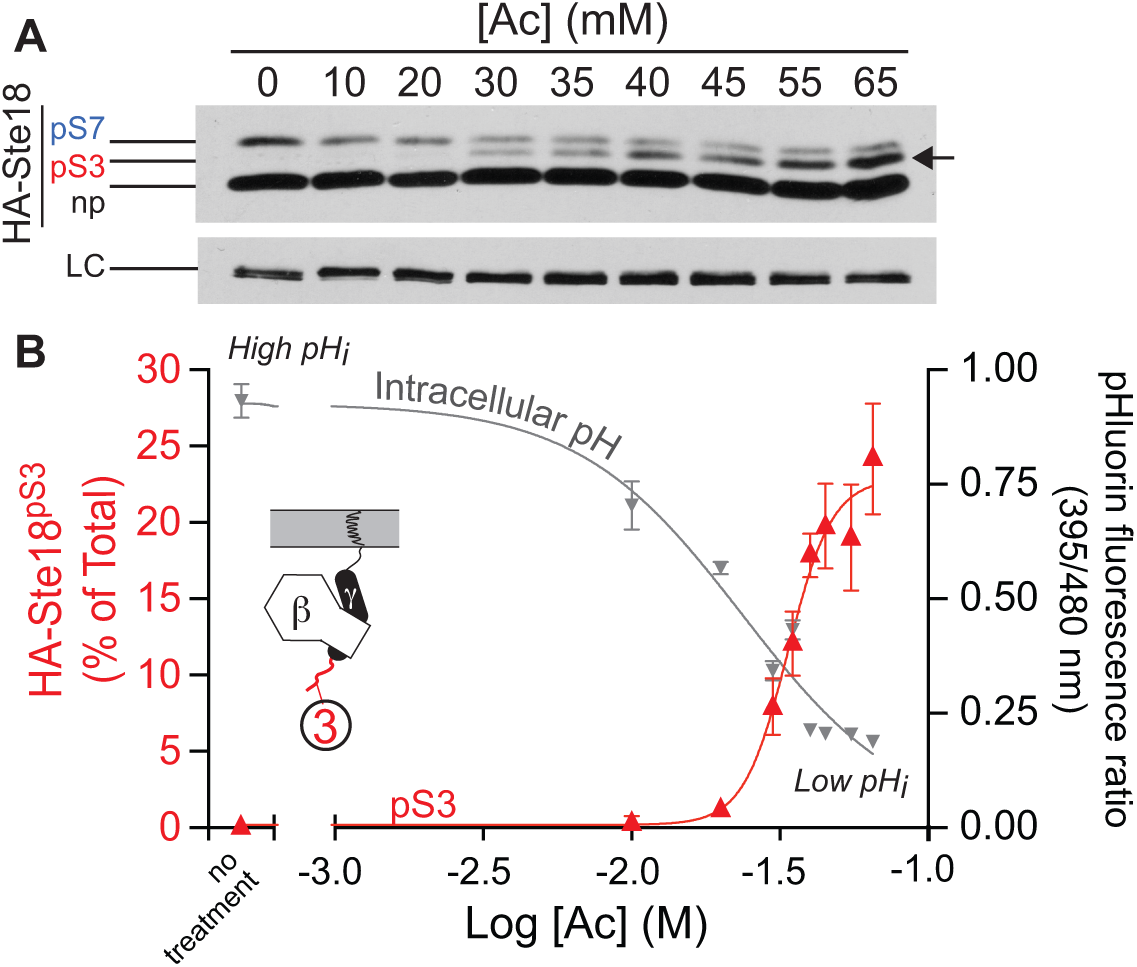
Ste18^pS3^ is a pH indicator. (A) Representative immunoblot showing Ste18^Nt^ phosphorylation upon 15 minute exposure to increasing concentrations of acetic acid in the absence of buffer. (B) Relative abundance of pS3 (red) and the ratio of fluorescence intensities emitted upon excitation at maximum pHluorin excitation wavelengths (395 nm/480 nm) as the function of Ac concentration logarithm (grey). Cells were collected 15 minutes after exposure to trap the maximal phosphorylation state of Ste18. Ste18 phosphorylation at Ser-7 (pS7) or Ser-3 (pS3), and non-phosphorylated Ste18 (np). (LC) loading control corresponding to yeast GAPDH. Results are the mean ± SEM (n=4).

### Pkc1, but not other CWI pathway kinases, is required for maximal phosphorylation of Ste18^S3^

Adaptation of yeast to acid stress has been shown to proceed through the cell wall integrity (CWI) pathway controlled by Pkc1 kinase (*30*). More recently, Pkc1 activated by mechanical stress has also been shown to phosphorylate Ste5 within the RING domain that mediates interaction of the scaffold protein with Ste4 and Ste18 (*31*). While Ste18^S3^ is not contained within a Pkc1 consensus sequence (*32*), we hypothesized that other downstream kinases in the CWI may potentially phosphorylate the Gγ subunit in response to acid stress. We tested this by measuring acid stress-dependent S3 phosphorylation in CWI kinase gene deletion strains. Surprisingly, we observed that the percentage of Ste18 phosphorylated on S3 was significantly diminished (∼3-fold) in cells lacking *PKC1* compared to wild type cells (Figure 5A,B). In contrast, pS3 was unaffected in cells lacking genes for downstream kinases – *BCK1, MKK1/MKK2, or SLT2* – and was even slightly elevated in *mkk1*Δ/*mkk2*Δ cells, in which total Ste18 abundance was also elevated (Figure 5A,B left). As expected, basal phosphorylation at S7 was not significantly altered in any of the strains (Figure 5B). Both basal and acid stress-dependent activation of the terminal CWI pathway kinase, Slt2, were completely blocked in *pkc1*Δ cells, which confirmed that maximal S3 phosphorylation requires Pkc1, but not downstream kinases of the CWI pathway (Figure 5C). Together these results imply that maximal acid stress-dependent phosphorylation of S3 is partially dependent on CWI-pathway kinase, Pkc1, as well as other non-CWI kinase(s) (Figure 5D).

**Figure 5.**
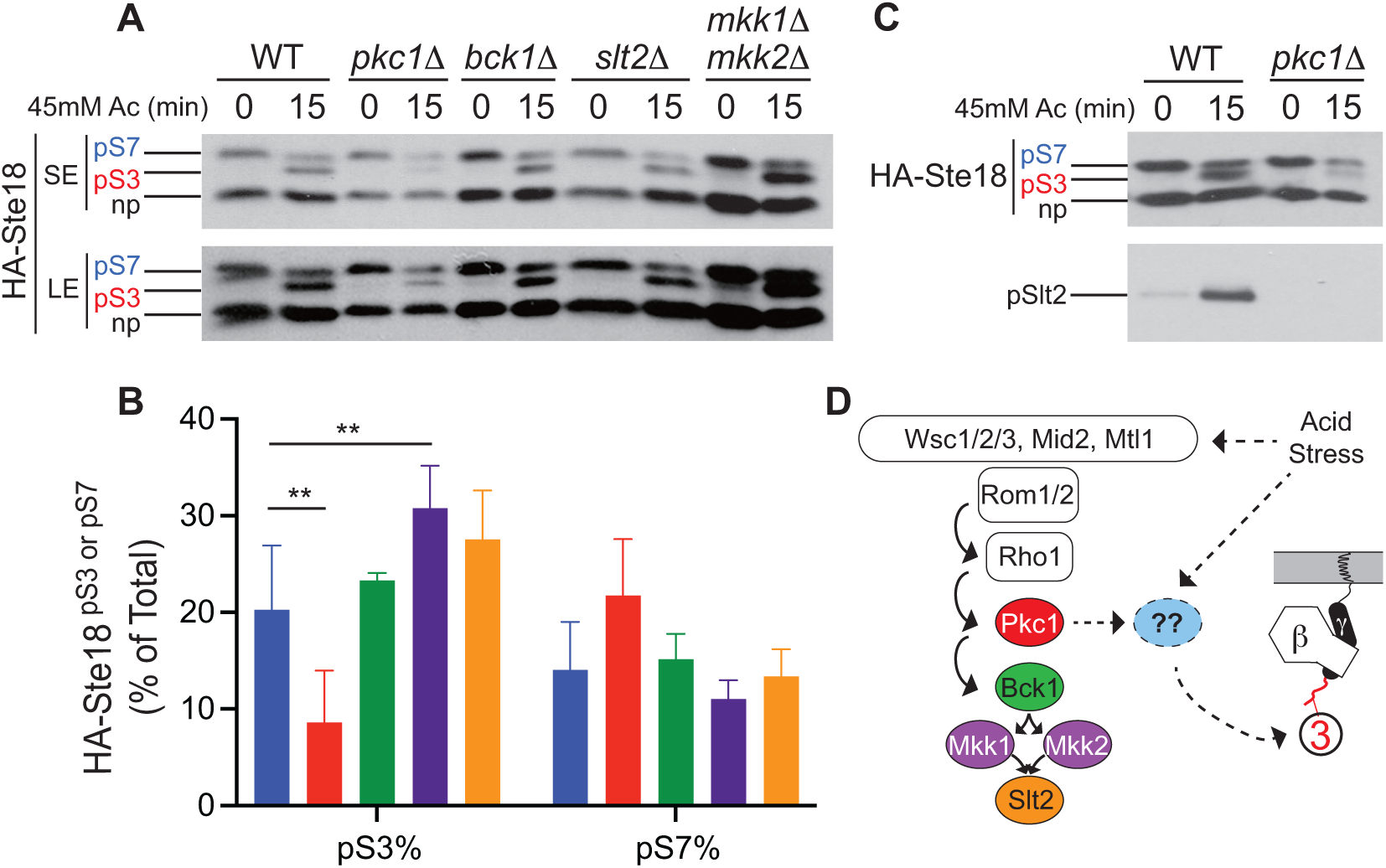
Pkc1, but not the CWI pathway, is necessary for maximal phosphorylation of Ste18^S3^ in response to acid stress. Wild type or single gene deletion mutant yeast strains were transformed with *pRS316-HA-STE18* (WT, *pkc1*Δ, *bck1*Δ, *slt2*Δ) or *pRS313-HA-STE18* (WT, *mkk1*Δ/*mkk2*Δ) under control of the *STE18* promoter/terminator followed by treatment with 45mM acetic acid without buffer for 0 or 15 minutes and SDS-PAGE and immunoblotting with anti-HA antibody. (A) Representative immunoblot of HA-Ste18 in wild type or different CWI pathway kinase gene deletion strains. (SE) Short exposure; (LE) Long exposure. (B) Quantification of pS3 and pS7 percentage after 15 minute exposure. Error bars represent the mean +/- standard deviation across 4-13 biological and analytical replicates (*see experimental procedures*). Statistical significance determined relative to WT using 2-way ANOVA with Tukey’s test for post-hoc multiple-comparisons analysis. (C) Representative immunoblot of activated Slt2 (pSlt2) in wild type and *pkc1*Δ cells in response to 0 or 15 minute exposure to 45mM acetic acid. (D) Schematic diagram of the CWI pathway showing potential interactions between acetic acid stress, the CWI pathway, an unknown kinase (blue circle), and Ste18.

### Ste18^pS3^ and Ste18^pS7^ are interactive

Intra-protein PTM interactions, in which the modification of one site in a protein affects the modification of another, often nearby, modification site in the same protein, are common in well-characterized regulatory IDRs (*8, 33–36*). To investigate whether S3 and S7 phosphosites are interactive on Ste18^Nt^, we quantified phosphorylation at each site in the presence and absence of phospho-nullifying or mimicking mutations at the neighboring site. We found that S3A had no impact on phosphorylation of S7 in response to either acid stress or pheromone stimulation (Figure 6A,B, S3). In stark contrast, the S3E mutation significantly dampened both pheromone-dependent, and acid stress-dependent phosphorylation at S7 by more than 50% compared to wild type cells (Figures 6C,D & S2). This effect on pS7 was uniform across all time points within each condition, suggesting that the efficiency, but not the dynamics of S7 phosphorylation, are sensitive to phosphorylation at S3. Thus, phosphorylation at S3 is likely to negatively impact phosphorylation at S7.

**Figure 6.**
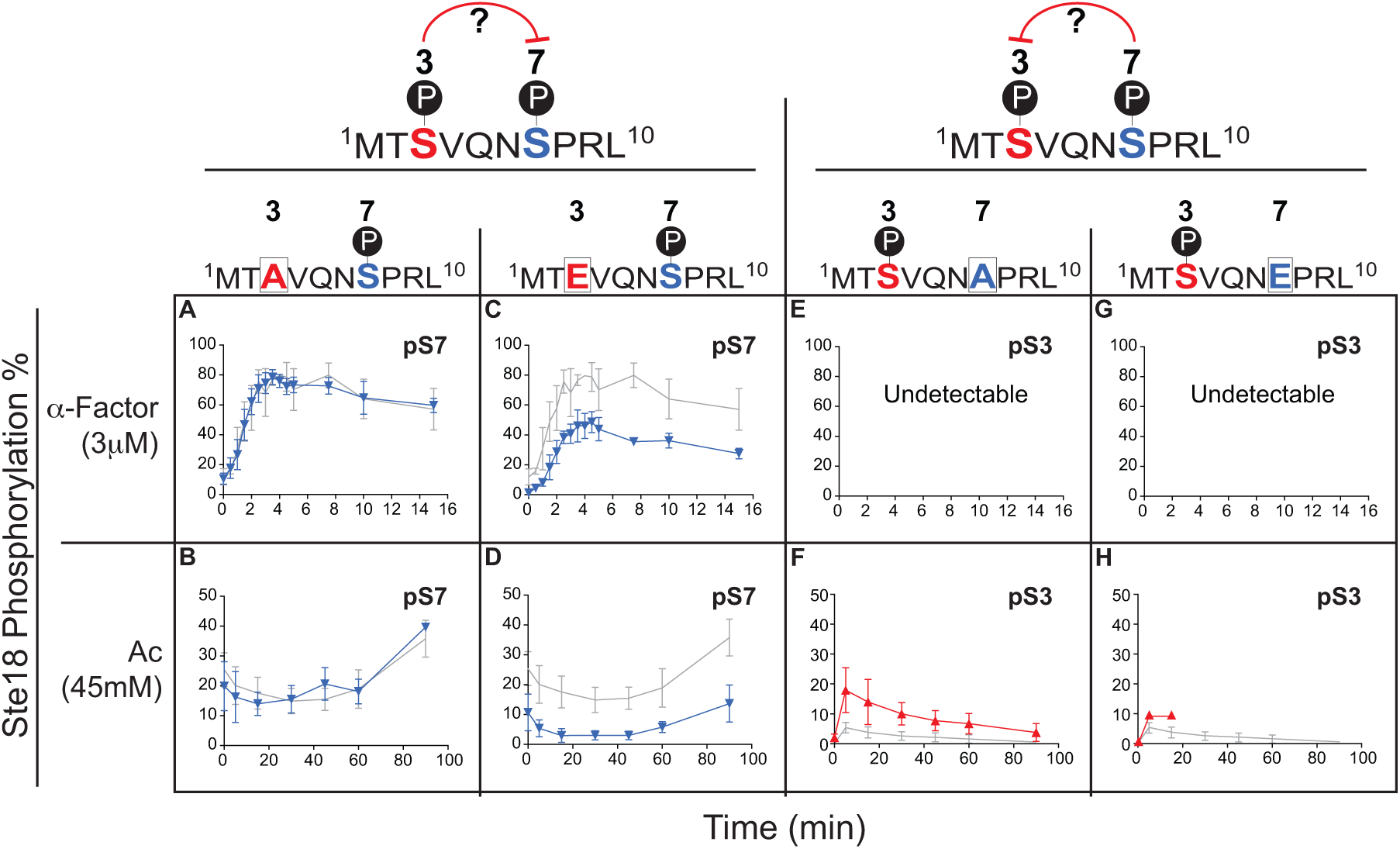
Site-specific substitution reveals interaction between Ste18^pS3^ and Ste18^pS7^. Ste18 pS3 and pS7 were measured by quantitative immunoblotting in yeast endogenously expressing wild-type or Ste18 phosphosite mutants exposed to 3μM pheromone or 45mM acetic acid in buffered media for 90 or 15 minutes, respectively. (A-D) Quantification of Ste18 pS7 with respect to S3A, S3E in pheromone or acid stress as indicated (blue). (E-H) Quantification of Ste18 pS3 with respect to S7A, S7E in pheromone or acid stress as indicated (red). Wild type responses (mean ± SD) across all experiments for each respective phosphosite in Ste18 wherein S3 and S7 are undisrupted (grey lines; n=6). Mutant results are the mean ± SD (n=4 to 6) for all but panel H pS3 (n=1). *See supplementary figure S2 for representative immunoblot images*.

Reciprocal testing the effects of S7 mutations on pS3 further revealed that pS3 and pS7 are differentially sensitive to mutation at the neighboring phosphosite. pS3 was not observed in pheromone treated S7A or S7E cells (Figure 6E,G), but did show a marked 3-fold increase from ∼6 to ∼18% in response to acid stress in S7A cells (Figure 6F), suggesting the phosphorylation of S7 negatively impacts phosphorylation at S3. This was confirmed in S7E cells, in which acid-dependent phosphorylation on S3 was diminished by ∼2-fold relative to S7A cells (compare Figures 6F,H), despite being higher than phosphorylation observed in wild type cells (Figure 6H). Taken together, these data provide strong evidence of a mutually exclusive interaction between pS3 and pS7 in Ste18, whereby phosphorylation on one site dampens phosphorylation on the other site.

### Phosphorylation alters the structure of Ste18^Nt^ *in vitro*

Recent evidence indicates that multi-site phosphorylation in an IDR can in some cases produce intrinsic disorder-to-order transitions (*37–39*). To asses this possibility for Gγ subunit N-terminal IDRs, we analyzed the effects of site-specific phosphorylation on the structure of the Ste18 N-terminus (residues 1-13) by molecular dynamics (MD) simulations coupled with circular dichroism (CD) spectroscopy. MD analysis of each 13-mer peptide isoform suggested a mild tendency for the IDR to adopt an alpha helical structure in the vicinity of the phosphosite - occurring near the extreme N-terminus when S3 is phosphorylated and further C-terminal when S7 is phosphorylated (Figure 7A,B & S3). Comparatively, the secondary structure propensity of the non-phosphorylated (NP) and fully phosphorylated (pS3pS7) peptides was considerably lower (Figure 7C,D & S3).

**Figure 7.**
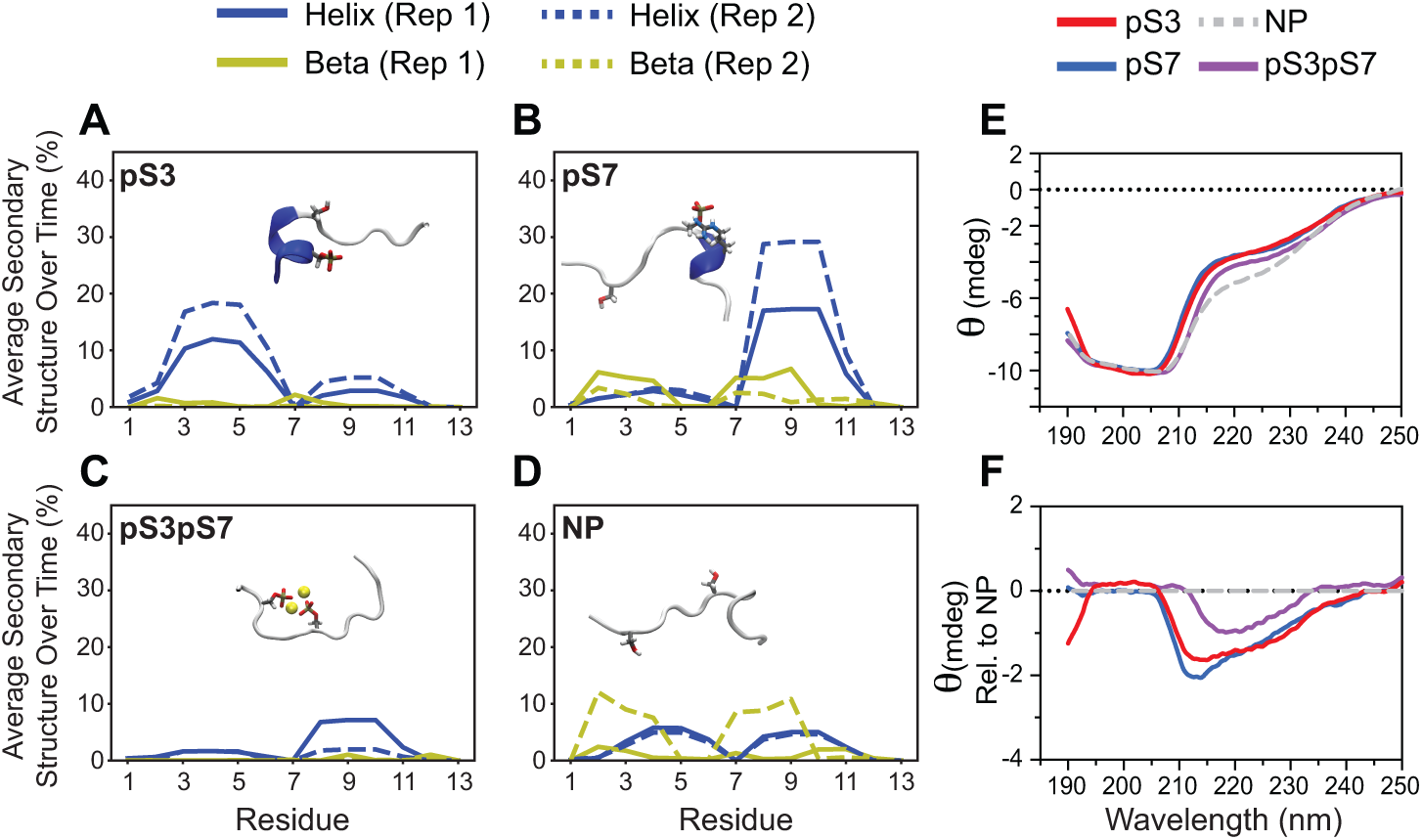
Phosphorylation promotes changes in Ste18^Nt^ IDR structure. (A-D) Two independent molecular dynamics simulations for synthetic pS3, pS7, pS3pS7, and non-phosphorylated (NP) peptides. Plots show the average secondary structure percentage (alpha helix or beta hairpin) relative to the amino acid position in each peptide (*see supplementary figure S3 for independent time traces*). (insets) Representative structures of the respective Ste18^Nt^ peptides. (E) CD spectrum of synthetic non-phosphorylated (NP) and phosphorylated Ste18^Nt^ peptides. Peptides are N-terminally acetylated and C-terminally amidated modifications to prevent N and C-terminal charge interactions that would not be possible within the N-terminus of the protein. (F) Difference CD spectra from E showing deviations between each phosphopeptide relative to the non-phosphorylated form.

The general trends observed by MD simulation were corroborated by differential CD spectropolarimetry of synthetic phosphopeptides. First, random coil is the dominant feature of each peptide, regardless of phosphorylation state and consistent with the prominence of this characteristic across all MD simulations (Figure 7E & S3). Second, pS3 and pS7 peptides produce the largest spectral differences with the greatest overlap, displaying more secondary structural character relative to the non-phosphorylated form (Figure 7F & S3). Third, the doubly phosphorylated peptide (pS3pS7) produces the least prominent difference spectrum relative to the non-phosphorylated peptide (Figure 7F) and consistent with the relative rarity of alpha or beta secondary structural character in the MD simulation of these peptides (Figure 7C,D). We concluded that the N-terminal IDR of Ste18 is innately intrinsically disordered and that site specific, but not combined phosphorylation, promotes weak but detectable secondary structure within this region.

### Combinatorial phosphorylation on Ste18^Nt^ tunes MAPK activation *in vivo*

Combinatorial protein modification is defined in one sense by the occurrence of distinct spatial combinations of protein modification that yield distinct functional outcomes (*8*). Phosphorylation of Ste18^Nt^ controls Fus3 activation rate and amplitude. Phospho-nullifying mutations enhance the activation rate and amplitude of Fus3 while phosphomimic mutations do the opposite (*18*). Moreover, these effects of the Gγ tail become hyper-exaggerated in the presence of Ste5^ND^ (*18*) – a docking mutant of the Gβγ effector Ste5 that prevents its binding to, and regulation by, Fus3 (*40*). Therefore, we used Fus3 activation kinetics in cells expressing Ste5^WT^ or Ste5^ND^ to report on the functional impact of combinatorial phosphorylation in Ste18^Nt^. Since pheromone does not induce S3 phosphorylation, we combined glutamate substitutions at position 3 (S3E) to mimic the effects of acid-induced S3 phosphorylation with alanine substitutions at position 7 to nullify naturally occurring S7 phosphorylation inherent to the pheromone response, resulting in four distinct protein isoforms: S3/S7 (*aka wild type*), S3/A7, E3/S7, and E3/A7. This approach minimized the potential for over-engineering the protein so as to capture realistic effects of combinatorial phosphorylation (i.e. serine rather than alanine is maintained as the natural residue for position); and also circumvented treatment of cells with acetic acid, which broadly effects many proteins in addition to Ste18 that could preclude any ability to decipher Ste18-specific effects on G protein signaling. Each Gγ isoform was expressed endogenously in either Ste5^WT^ or Ste5^ND^ cells, which were then stimulated with pheromone to monitor Ste18 phosphorylation and MAPK activation. We found that pheromone-induced pS7 exhibited identical behavior between Ste5^WT^ and Ste5^ND^ cells, confirming that disruption of the Fus3 regulatory site on Ste5 has no effect on Ste18 phosphorylation at S7 (Figure 8A,B & S4), and consistent with previously reported data (*18*). Likewise, we found no difference in S3E-mediated inhibition of S7 phosphorylation between Ste5^WT^ and Ste5^ND^ strains (Figure 8A,B & S4).

**Figure 8.**
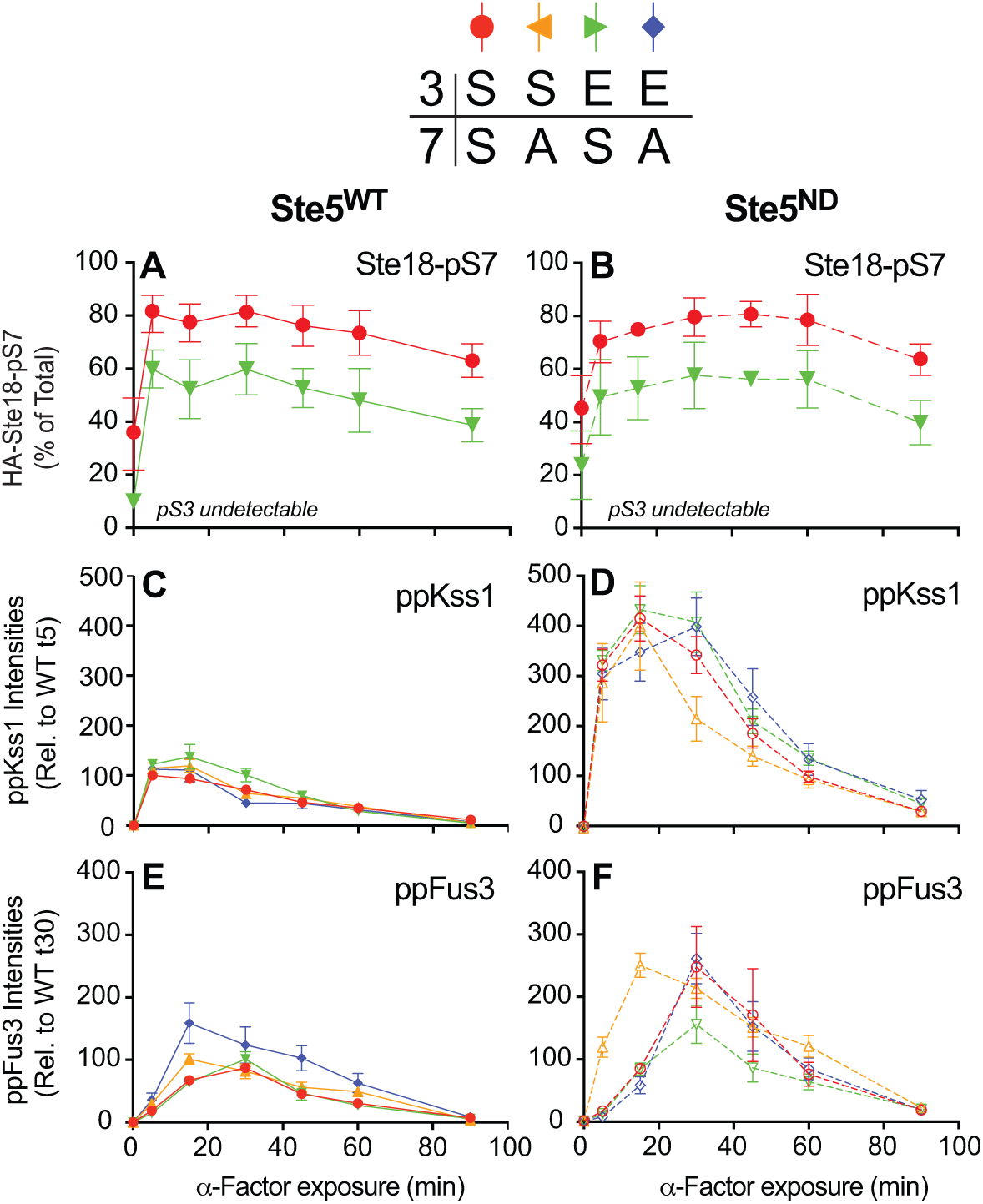
Combinatorial phosphorylation on Ste18^pS3^ and Ste18^pS7^ controls the activation rate and amplitude of MAPK Fus3. Fus3 activation is a hypersensitive diagnostic for the effects of Ste18^Nt^ phosphorylation when combined with Ste5 mutants lacking a functional Fus3 binding domain (Ste5^ND^) (*18*), and was used to evaluate combinatorial phosphorylation states. Yeast strains endogenously expressing Ste5^WT^ or Ste5^ND^ and one of four Ste18 phosphosite isoforms (S3/S7, S3/A7, E3/S7, E3/A7) were exposed to pheromone for 90 minutes with sampling for immunoblot analysis of HA-Ste18, activated Kss1 (ppKss1) and activated Fus3 (ppFus3), as indicated. (A,B) Relative abundance of pS7 in Ste5^WT^ (A) and Ste5^ND^ (B) cells. (C,D) Time-resolved activation of Kss1 in Ste5^WT^ (C) and Ste5^ND^ (D) cells. (E,F) Time-resolved activation of Fus3 in Ste5^WT^ (E) and Ste5^ND^ (F) cells. Results are the mean ± SEM (n=8). *See figure S4 for representative immunoblot images*.

Next, we compared activation of the pheromone-responsive MAPKs in each strain. Activation of the yeast mating pathway triggers phosphorylation of Fus3 and also Kss1, which can both be monitored simultaneously. Overall, Kss1 activation was relatively unaffected by Ste18 phosphosite mutants in either the Ste5^WT^ or Ste5^ND^ background, despite the large increase in its overall activation in Ste5^ND^ cells that has been documented previously (Figure 8C,D & S4) (*41*). In contrast, we found that Fus3 activation kinetics are dependent on the phosphorylation state of Ste18. In Ste5^WT^ cells, Fus3 activation is mildly sensitive to the S7A mutation, resulting in faster peak activation that occurred at 15 minutes rather than the typical 30 minute peak in Ste5^WT^ cells (Figure 8E & S4). In Ste5^ND^ cells, Fus3 activation was more pronounced and revealed functional distinctions between the different Ste18 phosphosite mutants. Ste5^ND^ cells expressing wild type Ste18 (S3/S7) produce a characteristic peak of Fus3 activation at 30 minutes but with ∼2.5-fold greater total signal than occurs for Ste5^WT^ cells (Figure 8F). S7A (i.e. *S3/A7*), which prevents all phosphorylation in response to pheromone, does not change Fus3 activation intensity but does shorten peak activation time by 2-fold to 15 minutes. In contrast, E3/S7, which models a form of Ste18 that is maximally phosphorylated at both S3 and S7, represses Fus3 activation without altering the rate of peak activation in the Ste5^ND^ background. Lastly, we analyzed E3/A7, which models a form of Ste18 that is only phosphorylated at position 3 but not at position 7 (Figure 8F). Cells expressing this form of Ste18 exhibited wild type (i.e. *S3/S7*) behavior when combined with Ste5^ND^. Taken together, these data suggest that phosphorylation in the N-terminal IDR of a G protein γ subunit, and in Ste18 specifically, can have distinctive structural and functional effects that govern G protein signaling outcomes.

## Discussion

We have presented here the first evidence that the intrinsically disordered N-terminal tails of G protein γ subunits can undergo stimulus-dependent, combinatorial, multi-site phosphorylation that alters their inherent structure and function. We have characterized in detail the site-specific N-terminal phosphorylation of Ste18, the single canonical Gγ subunit required for pheromone-dependent G protein signaling in yeast. Our results show that Ste18^Nt^ is differentially phosphorylated at two unique serine residues, S3 and S7, in response to a range of stimuli including pheromone-dependent GPCR activation, osmotic stress, glucose stress, and pH stress. Phosphorylation of S3, in particular, is uniquely sensitive to pH stress and promotes a state in which both sites are phosphorylated simultaneously. We exploited this fact to test the hypothesis that Gγ subunits are among the short list of proteins, such as histones and GPCRs, that undergo functional combinatorial modification within their terminal IDRs. Through a combination of *in vivo* and *in vitro* approaches we provide several pieces of evidence in support of this hypothesis,

showing specifically that: one, phosphorylation between S3 and S7 is likely interactive such that phosphomimicry at one site negatively affects phosphorylation on the other site; two, phosphorylation induces subtle yet readily detectable structural changes in the N-terminal IDR *in vitro*; three, the structural impact of single versus double site phosphorylation are distinct; and four, that different patterns of phospho-mimicry or phosphorylation on S3 and S7 result in distinct MAPK activation profiles *in vivo*. The sum of these data provides strong support for the hypothesis that Gγ N-terminal IDRs are multi-conditional targets of cell stress-mediated combinatorial phosphorylation that governs G protein signaling.

### Phosphorylation, not structural change, likely dominates the regulatory effects of Ste18^Nt^

Generally speaking, the functional effect of any PTM can be mediated through two possibilities. A modification may elicit a change in structure and this structural change alters protein function (through altered tertiary or quaternary protein structure). Alternatively, a modification may itself disrupt or promote function by creating a new surface with which proteins may interact or by disrupting an existing surface necessary for interaction. IDRs are inherently disordered, and as such, their modification is often suspected to mediate functional effects through the second of these two options. However, emerging evidence suggests that PTMs on IDRs can elicit disorder-to-order transitions, such as the formation of phosphorylation-stabilized beta sheets (*39*). Our work is the first to investigate the structural effects of phosphorylation on an N-terminal IDR within a canonical Gγ subunit. The results suggest that, despite some minor changes in observable secondary structure, the intrinsic disorder of these regions is largely persistent, regardless of phosphorylation state (Figure 7). Consequently, we did not find a strong correspondence between the structural and functional effects of phosphorylation in the Ste18^Nt^ (compare Figures 7 & 8). Therefore, our data suggest that phosphorylation, more so than structural changes induced by phosphorylation, mediate the regulatory properties of Gγ N-terminal tails.

### Multiple phosphosites in Ste18^Nt^ relay crosstalk from a range of cellular stress pathways

Previous work had shown that pheromone and MAPK-dependent feedback phosphorylation of Ste18^Nt^ negatively regulates Ste5-dependent plasma membrane association and phospho-activation of Fus3 phosphorylation (*18*). Our work has expanded significantly on this to show that the Ste18^Nt^ phosphorylation occurs at multiple sites and in response to a variety of cellular stress stimuli including osmotic, glucose, and pH-dependent stress. This discovery is highly consistent with the role of Ste18^Nt^ phosphorylation as a negative regulator of MAPK signaling and suggests that in yeast, phosphorylation of the Gγ subunit is one of many ways in which negative feedback can be relayed to the mating response. Indeed, we confirm that under hyperactivating conditions facilitated by mutations in the Gβγ effector Ste5, activation of the primary mating-specific MAPK Fus3, occurs unhindered in the absence of Ste18^Nt^ phosphorylation (see 3S/7A), is partially repressed in response to single site phosphorylation alone (3S/pS7 and 3E/7A), and is maximally repressed in response to dual site phosphorylation (see 3E/pS7) (Figure 8F). Since phosphorylation of S3 and S7 are distinct in their conditional dependence (where pS3 is responsive to low glucose or acid stress and pS7 is responsive to pheromone stimulation or osmotic stress), they function like cellular stress indicators that relay crosstalk from a range of stress pathways and broaden the capacity of Gβγ-dependent signaling to be modulated in response. Furthermore, phosphorylation is dynamic and responsive whereby it is maintained only for the duration of the stress or until the cell has equilibrated to the stress, which can be seen in the short pulse of pS7 observed under osmotic stress (Figure 1B), the elongated response to acid stress (Figure 3C), and the temporary response to acid in buffered culture media (Figure 3F).

While our data provides the first evidence that such crosstalk can govern G protein signaling through Gβγ subunits, new questions also arise from this work. For example, the stoichiometry of Ste18 phosphorylation in response to cellular stress (5-20%) is rather low compared to phosphorylation in response to pheromone (∼80%), which could suggest that Ste18^Nt^ may not be a primary or perfect target for kinases activated under the stresses we’ve tested, that there are strict cellular localization constraints for crosstalk to occur efficiently under each stress, or that we have simply gleaned the potential for stress crosstalk through Ste18^Nt^ but have not yet identified the *linchpin* stressor or stress condition that would promote high stoichiometry phosphorylation and regulation. Indeed, our first glimpse into Ste18 phosphorylation came from work done in osmotic stress, only to find later that pheromone-dependent phosphorylation was more predominant (*18*). While answering these questions will be of potential benefit to our understanding of the mechanism, the experimental results reported here have shown what the ultimate signaling outcome of stoichiometric site-specific phosphorylation would be.

### Multiple conditions – multiple sites – multiple kinases

The kinases that phosphorylate S3 and S7 in Ste18^Nt^ are different, not only due to the uniqueness of each site in the protein, but also due to the conditions under which they become phosphorylated. Previous evidence suggests that Fus3 is responsible for phosphorylation of S7 in response to pheromone-dependent GPCR activation (*18*). However, the fact that Fus3 is not activated by osmotic stress indicates that other S7 kinases must also exist. Furthermore, basal phosphorylation, which is unique in that it is stimulus independent, suggests the involvement of a third S7 kinase. Thus, beyond the fact that multiple phosphosites exist in Ste18^Nt^, condition-specific phosphorylation patterns suggest that a site can be targeted by multiple kinases relaying crosstalk through the same residue.

In contrast to S7, S3 does not fit within a well-defined kinase consensus sequence and is difficult to predict by sequence alone (*42*). Surprisingly, the CWI pathway, which is activated by acid stress, is not essential for phosphorylation at this site since we were able to observe S3 phosphorylation even in the absence of the primary CWI kinase, Pkc1 (Figure 5). However, the significant decrease in pS3 observed in *pkc1*Δ cells does suggest that Pkc1, but not downstream kinases, is necessary for maximal phosphorylation. The consensus sequence for Pkc1 phosphosites, generated by hundreds of different substrates (*32*), suggests that S3 is not a direct target of the kinase, but rather another kinase whose action on Ste18 is Pkc1-dependent (Figure 5D). In any case, we have shown that S3 phosphorylation functions as a quantitative indicator of intracellular pH (Figure 4), and in view of the fact that the complete CWI pathway is not essential for this suggests that pH itself may activate the responsible kinase. Indeed, emerging evidence that protons can function as second messengers in G protein signaling pathways (*27, 43, 44*), provides support for this hypothesis. Nonetheless, identifying the complete suite of Ste18 kinase(s) could shed further light on the mechanism underlying pH-dependent phosphorylation at Ste18^S3^ and the identity of the pH sensor(s) responsible for this effect.

### Interdependence of Gγ phosphorylation and Gβγ effectors

While we have shown the effect of Ste18 phosphorylation is significant in yeast, it is also not completely independent of the Gβγ effector Ste5. Indeed, we confirmed that the effects of Gγ phosphorylation are most prominent when coupled with the Fus3 non-docking mutant of Ste5 (Ste5^ND^), which cannot be phospho-regulated by Fus3 (Figure 8F). Indeed, previous work has shown that genetic mutations on Ste18^Nt^ and Ste5 synergize to control MAPK signaling by greater than the sum of their individual effects (*18*). More recently, Pkc1 was shown to phosphorylate Ste5 within the Ste5^RING^ domain that mediates interaction of the scaffold protein with Ste4/Ste18 (*31*). Thus, co-regulation of both Gβγ subunits and their effectors may play complementary roles in the modulation of Gβγ signaling. In yeast, additional Ste4/Ste18 effectors (Far1, Ste20, Cdc24) are also key players in the Gβγ-dependent pheromone response and whether they, like Ste5, synergize with phosphorylated Ste18 remains unknown. In humans there are 5 Gβ and 12 Gγ subunits, which can form a wide variety of Gβγ dimers that interact directly with several different effectors (*45*). Many of these Gγ subunits are phosphorylated in their N-terminal IDRs (*19, 46*), but have not been studied to elucidate the functional importance of these sites – especially in terms of their dependence on effector modification state. Thus, continued work in this area can expand the existing paradigm of Gβγ signaling not only in yeast but also in other organisms where there is growing evidence to support the hypothesis that N-terminal IDRs of Gγ subunits are combinatorially-regulated governors of G protein signaling.

## Experimental Procedures

### Yeast strains and plasmids

Unless specified, all strains used in this study are derived from the *BY4741* (*MATa leu2Δ met15Δ his3Δ ura3Δ*) background and are listed in the supplementary information (Table S1). All mutants were constructed using delitto perfetto mutagenesis (*47*) and verified by PCR amplification and dideoxy sequencing (Eurofins MWG Operon). Cells lacking *PKC1* (*pkc1*Δ; *DL376*) and the isogenic wild type strain (*DL100(1783)*) (*48*), were graciously provided by Dr. David Levin. To construct *pRS313*- and *pRS316-HA-STE18*, primers ZNT433 (5’TGGAGCTCCACCGCGGTGGCGGCCGCTAGGGCGCGACACGTCTAAA3’) and ZNT434 (5’TCGAATTCCTGCAGCCCGGGGGATCCCAGCGAGATTTATTTCGAAA3’) were used to amplify wild type *HA-STE18* including its endogenous promoter (−510bp) and terminator (+467bp) from the *YMT235* strain genome. The PCR product was cloned into pRS313 or pRS316 plasmids digested with *NotI* and *BamHI* restriction sites using the sequence and ligation independent cloning (SLIC) method (*49*).

### Media and growth conditions

Depending on treatment conditions, yeast strains were grown in either YPD growth medium (Yeast Extract, Peptone, 2% dextrose media) or synthetic complete (SC) (or SC drop-out) medium with 2% dextrose. When performing experiments in buffered conditions SC medium with 2% dextrose was buffered by addition of 50mM dibasic potassium phosphate, 50mM dibasic sodium succinate and adjustment to pH 5.0 by addition of HCl (*27*).

Cells were grown at 30°C to mid log phase (OD_600_ 0.75–0.85) followed by treatment as indicated for the specific experiment (see treatments below). After treating cells for the desired period of time, cell growth was stopped by adding 5% trichloroacetic acid (TCA) after which cells were immediately harvested by centrifugation at 3724 × g in Allegra X-14R Beckman Coulter Centrifuge at 4°C, washed with ice-cold Milli-Q water and frozen at -80°C.

Specific treatment conditions were as follows: *GPCR activation* – Cells were treated with α-factor peptide hormone (Genscript) at 3μM final concentration. *Osmotic Stress –* Cells were collected by centrifugation and re-suspended in the same medium supplemented with 0.75 M KCl for the desired amount of time, as described previously (*50*). *Glucose Stress –* Cells were collected by centrifugation, washed with the same medium lacking glucose and re-suspended in glucose free synthetic media supplemented with the desired amount of glucose for the desired time (*27*). *Acetic acid stress* – Cells were grown in buffered or unbuffered SC medium containing 2% dextrose followed by addition of acetic acid at 45mM final concentration unless otherwise noted (*27*).

### Cell lysis and western blotting

Cell pellets were subjected to glass bead lysis in the presence of TCA buffer as per the standardized protocol described previously (*51*). The protein concentration was measured using a DC Protein Assay (Bio-Rad). Protein extracts were separated by SDS-PAGE with either 7.5% or 12.5% acrylamide. Primary antibodies and dilutions used for immunoblotting included: phosphorylated Slt2, Kss1, and Fus3 (Phospho-p44/42 MAPK) (Cell Signaling Technologies, Cat #9101) (1:500); hemagglutinin antigen epitope (HA) (Cell Signaling Technologies, Cat #3724) (1:3000); glucose-6-phosphate dehydrogenase (loading control; LC) (Sigma-Aldrich, Cat #A9521) (1:50,000). In all cases, a horseradish peroxidase (HRP)-conjugated secondary antibody (goat-anti-rabbit) (Bio-Rad, Cat # 1705046) (1:5000) was used for detection. The signal was detected using enhanced chemiluminescence (ECL) reagent (PerkinElmer catalog #NEL 104001EA), developed on film E3018 HyBlot CL Autoradiography Film (Denville Scientific). In all cases, several exposures were taken and only those exposures for which all signals were below saturation were used for quantification. Quantification was achieved by hi-resolution scanning of appropriate films followed by image densitometry using ImageJ software to quantify signal intensities (*52*).

### Analysis of Ste18 phosphorylation in CWI kinase deletion strains

Analysis of Ste18 phosphorylation in kinase deletion strains was accomplished by transformation of each strain with *pRS316-HA-STE18 (WT, pkc1*Δ, *bck1*Δ, *slt2*Δ) or *pRS313-HA-STE18 (WT, mkk1*Δ*/mkk2*Δ). Cells were then grown in SC-URA or SC-HIS medium and treated with acetic acid as described above for up to 15 minutes. For *pkc1*Δ and *mkk1*Δ/*mkk2*Δ strains, the media was supplemented with 1M sorbitol to facilitate growth. Each transformed strain was grown in triplicate (biological replicates) and analyzed first with respect to the isogenic wild type strain for the specific mutant on a single blot. Each strain replicate was then re-analyzed on the same gel with the other deletion strains and wild type strain *(DL100(1783))* (analytical replicates). Ste18 phosphorylation percentages calculated from both biological and analytical replicates were included in the quantitative analysis.

### Phosphatase Assay

Cells were grown as described earlier in a non-buffered synthetic media and treated with 45mM acetic acid for 15 minutes followed by addition of TCA, cell harvesting by centrifugation and freezing at -80C. Cells were subjected to glass bead lysis in the presence of 1x phosphatase buffer mix (1x PMP buffer (New England Biolabs), 1x MnCl_2_, and 1x EDTA-free protease inhibitors (Roche)), as described previously (*18*). Lysates were clarified by centrifugation, the supernatant split, and one half treated with lambda protein phosphatase enzyme (New England Biolabs, P0753S) to the final concentration of 2.25 U/μl for 30 minutes at 30C. The other half incubated in the same manner without phosphatase treatment to serve as a control. 6x SDS loading dye was added to all reactions and boiled to stop the reaction, followed by SDS-PAGE and western blot analysis.

### Intracellular pH measurement

For intracellular pH (pH_i_) measurement, yeast cells were transformed with pZR4.1 plasmid encoding pHluorin under the *TEF1* constitutive promoter (*53*), and pH_i_ was measured as described previously (*27*). To create the calibration curves (Figure S5), mid-log phase cells expressing pHlourin were permeabilized with digitonin at 100 μg/ml final concentration (Sigma-Aldrich, Cat #D141) for 10 minutes at 30C. Permeabilized cells were washed with PBS (25mM potassium phosphate, 100mM KCl) (pH 7.0) followed by resuspension in PBS ranging from pH 5.0 to pH 8.0 (pH 5, 5.5, 6, 6.5, 7, 7.5, 8). The ratio of pHluorin emission fluorescence intensity at 520 nm produced by excitation at 395 nm and 480 nm was measured using a Synergy H1 hybrid fluorescence plate reader (Biotek). For intracellular pH measurements, Prism software, v8 (GraphPad Software) was used to convert pHluorin fluorescence intensities emitted upon excitation at the two excitation wavelengths to pH values using the calibration curve.

### Circular dichroism spectropolarimetry

Commercial synthetic Ste18^Nt^ peptides were synthesized with N-terminal acetylation and C-terminal amidation and various combinations of S3 and S7 phosphorylation (or non-phosphorylated) and enriched to 95% purity (Genscript). Lyophilized peptides were reconstituted in deionized water at a concentration of 2.5 mg/ml and further diluted as necessary with 10mM potassium phosphate buffer (pH 7). Far-ultraviolet (190-250 nm) circular dichroism (CD) spectroscopy was performed on a Jasco J-815 CD spectrometer equipped with Peltier temperature control unit. Peptide samples were diluted to 0.25 mg/ml in 10mM potassium phosphate buffer (pH 7). A quartz cuvette with 1mm path length was used at 25C in this experiment. Measurements were carried out with 50 nm/min scan rate in 0.2 nm steps at 1 second response time and 1 nm bandwidth. Single CD spectra were averaged over 15 scans after buffer baseline correction.

### Molecular Dynamics

MD simulations were carried out with NAMD2.13 (*54*) and Amber16 (*55*). The CHARMM22* protein (*56*) and CHARMM36 nucleic acid force field (*57*) were used to describe the peptides, and TIP3P water model (*58*) was used to describe the solvent/ions. The initial structure of the Ste18 N-terminus (residues 1-13) was prepared using VMD and its plugin, Molefacture (*59*). All four phosphorylated states of the peptide were constructed, including phosphorylation at Ser-3 (pS3), Ser-7 (pS7), both Ser-3 and Ser-7 (pS3pS7), and non-phosphorylated (NP). Each peptide was placed in a water box, and ions were added to neutralize the system at a concentration of 150 nM NaCl. The final system size was around 20,300 atoms.

Each system was equilibrated at 400K and constant volume for 4 ns to randomize the starting conformation, and then cooled at 298 K and 1 atm for 2 ns. Next, two independent 4 μs production runs were performed for each system at 298 K and constant volume. The temperature was maintained using Langevin dynamics for all simulations, and the pressure was kept at 1 atm using the Langevin piston method when applied (*60*). The equilibration simulations were performed using a 2-fs time step. Hydrogen mass repartitioning (HMR) was employed for the production runs to reach a time step of 4 fs (*61, 62*). All simulations were performed under periodic boundary condition with a cutoff at 12 Å for electrostatic and Lennard-Jones interaction and a switching function beginning at 10 Å. Long-range electrostatic interactions were calculated using the particle-mesh Ewald (PME) method (*63*). System setup, analysis and visualization were performed using VMD.

## Supporting information

NassirToosi_SupplementalInfo_2020

## Acknowledgements

We would like to thank Dr. Nicholas V. Hud (Georgia Institute of Technology) for CD spectropolarimeter access, Dr. Amit Reddi (Georgia Institute of Technology) for the pHluorin plasmid, Dr. David Levin (Boston University) for *pkc1*Δ yeast strains, and Dr Kuntal Mukherjee and Parastoo Baradaran-Mashinchi for Ste18 constructs. Special thanks to Daniel Isom for critical review and discussion. This work was funded in part by National Institutes of Health grant R01-GM117400 to M.T, R01-GM123169 to J.C.G., and by the Southeast Center for Mathematics and Biology via grants NSF-DMS1764406 and Simons Foundation/SFARI-594594 to M.T. Computational resources were provided through the Extreme Science and Engineering Discovery Environment (XSEDE; TG-MCB130173), which is supported by NSF Grant Nation. This work also used the Hive cluster, which is supported by the National Science Foundation under grant number 1828187 and is managed by the Partnership for an Advanced Computing Environment (PACE) at the Georgia Institute of Technology.

