## Supplementary material for "Combinatorial phosphorylation modulates the structure and function of the G protein gamma subunit in yeast": NassirToosi_SupplementalInfo_2020

**Table S1.** List of yeast strains used in this study

**Figure S1.** Immunoblots for HA-Ste18$^{WT}$ and HA-Ste18$^{S3A,S7A}$ after exposure to 45mM acetic acid in buffered medium.

**Figure S2.** Representative immunoblots for HA-Ste18 and indicated phosphosite mutant proteins after exposure to 45mM acetic acid or 3μM pheromone.

**Figure S3.** Time resolved secondary structure propensity of Ste18 residues 1-13 simulated through molecular dynamics (*see materials and methods*).

**Figure S4.** Representative immunoblots for activated Kss1 (ppKss1), activated Fus3 (ppFus3), HA-Ste18$^{WT}$ and HA-Ste18 mutants after exposure to 3μM pheromone.

**Figure S5.** pHluorin calibration curves.

**Table S1.** List of yeast strains used in this study.

| Strain | Description | Source |
|---|---|---|
| *BY4741* | *MAT**a** his3Δ1 leu2Δ0 met15Δ0 ura3Δ0* | PMID 9483801 |
| *YMT235* | *BY4741 HA-STE18 WT* | PMID 26070665 |
| *YMT 469* | *BY4741 HA-STE18 T2A* | this study |
| *YMT 502* | *BY4741 HA-STE18 T2E* | this study |
| *YMT 470* | *BY4741 HA-STE18 S3A* | this study |
| *YMT 503* | *BY4741 HA-STE18 S3E* | this study |
| *YMT 471* | *BY4741 HA-STE18 S7A* | this study |
| *YMT 504* | *BY4741 HA-STE18 S7E* | this study |
| *YMT 590* | *BY4741 HA-STE18 S3E-S7A* | this study |
| *YMT 522* | *BY4741 HA-STE18 S3A,S7A* | this study |
| *YMT 491* | *BY4741 HA-STE18 WT STE5-ND (Q292A, I294A, Y295A, L307A, P310A, N315A)* | PMID 29719261 |
| *YMT 533* | *BY4741 HA-STE18 S7A STE5-ND* | this study |
| *YMT 561* | *BY4741 HA-STE18 S3E STE5-ND* | this study |
| *YMT 608* | *BY4741 HA-STE18 S3E,S7A STE5-ND* | this study |
| *DL100(1783)* | *MATa leu2-3,112 ura3-52 trpl-1 his4 can1'* | PMIDs 1740473, 19805511 |
| *DL376* | *MAT**a** leu2-3,112 ura3-52 trpl-1 his4 can1' pkc1Δ::LEU2* | PMID 1740473 |
| *bck1Δ* | *BY4741 bck1Δ::kanMX4* | PMID 9483801 |
| *mkk1Δmkk2Δ* | *BY4741 mkk1Δ::Kan-MX4 mkk2Δ::CORE(kanMX4 Kl-URA3)* | PMID 9483801; this study |
| *slt2Δ* | *BY4741 slt2Δ::kanMX4* | PMID 9483801 |

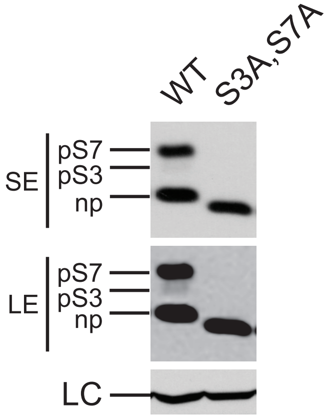


**Figure S1.** Immunoblots for HA-Ste18$^{WT}$ and HA-Ste18$^{S3A,S7A}$ after exposure to 45mM acetic acid in buffered medium. LC, GAPDH loading control.

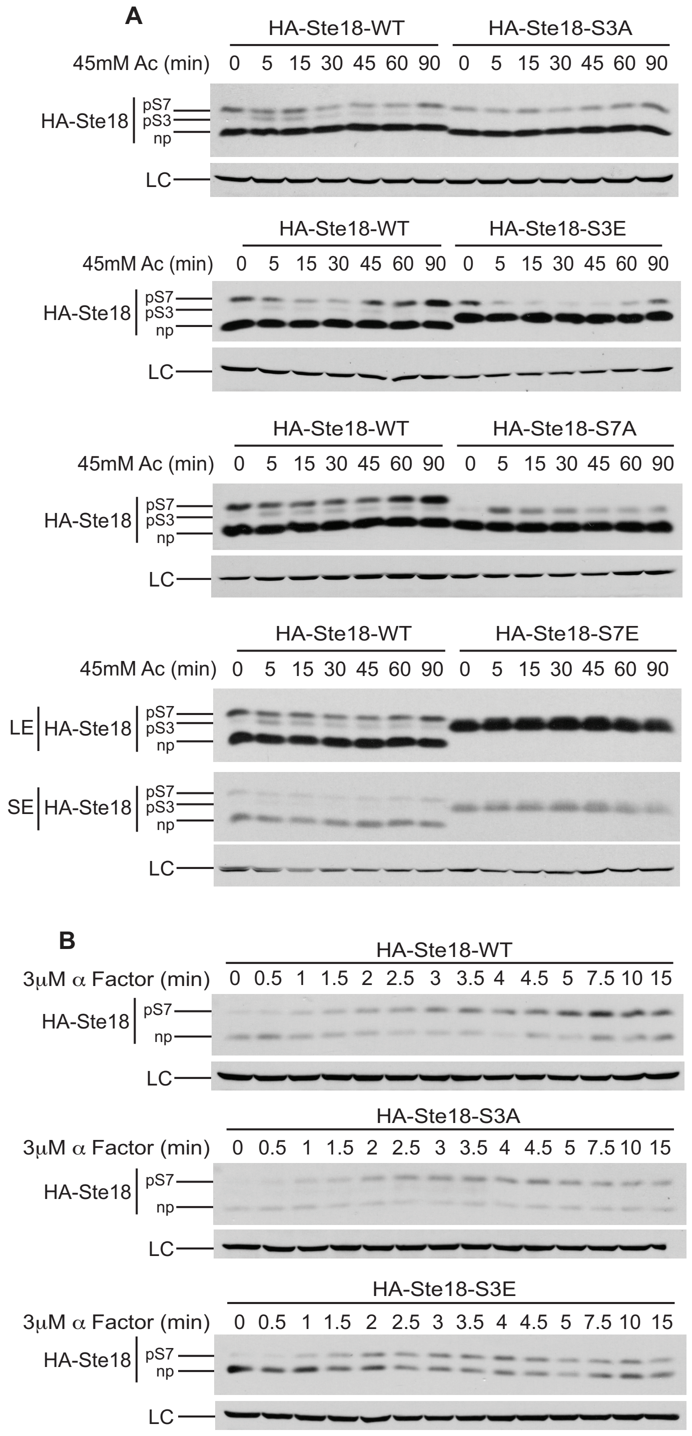


**Figure S2.** Representative immunoblots for HA-Ste18 and indicated phosphosite mutant proteins after exposure to 45mM acetic acid (A) or 3μM pheromone (B) for the indicated time. SE, short exposure; LE, long exposure; LC, GAPDH loading control.

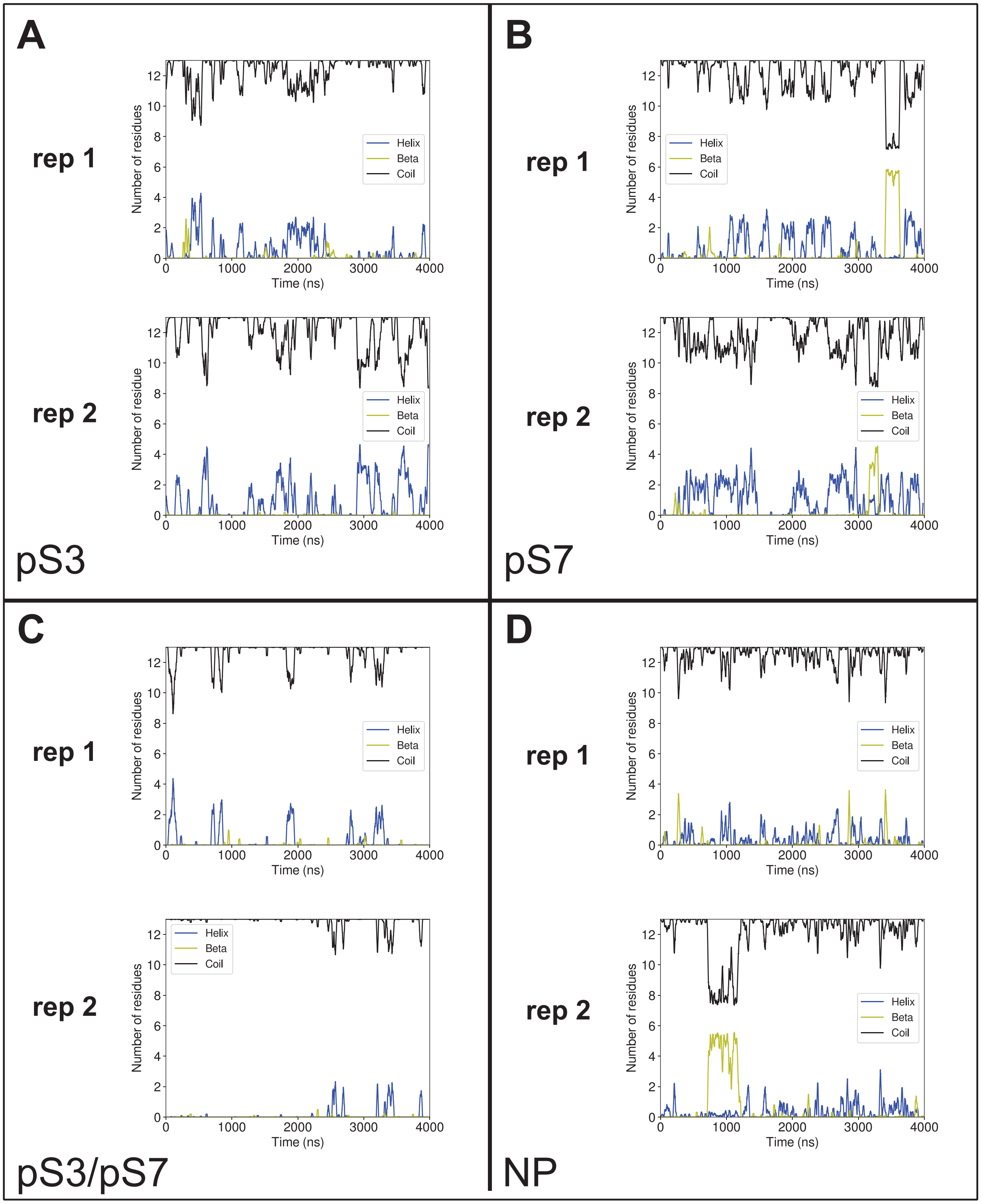


**Figure S3.** Time resolved secondary structure propensity of Ste18 residues 1-13 simulated through molecular dynamics (*see materials and methods*).

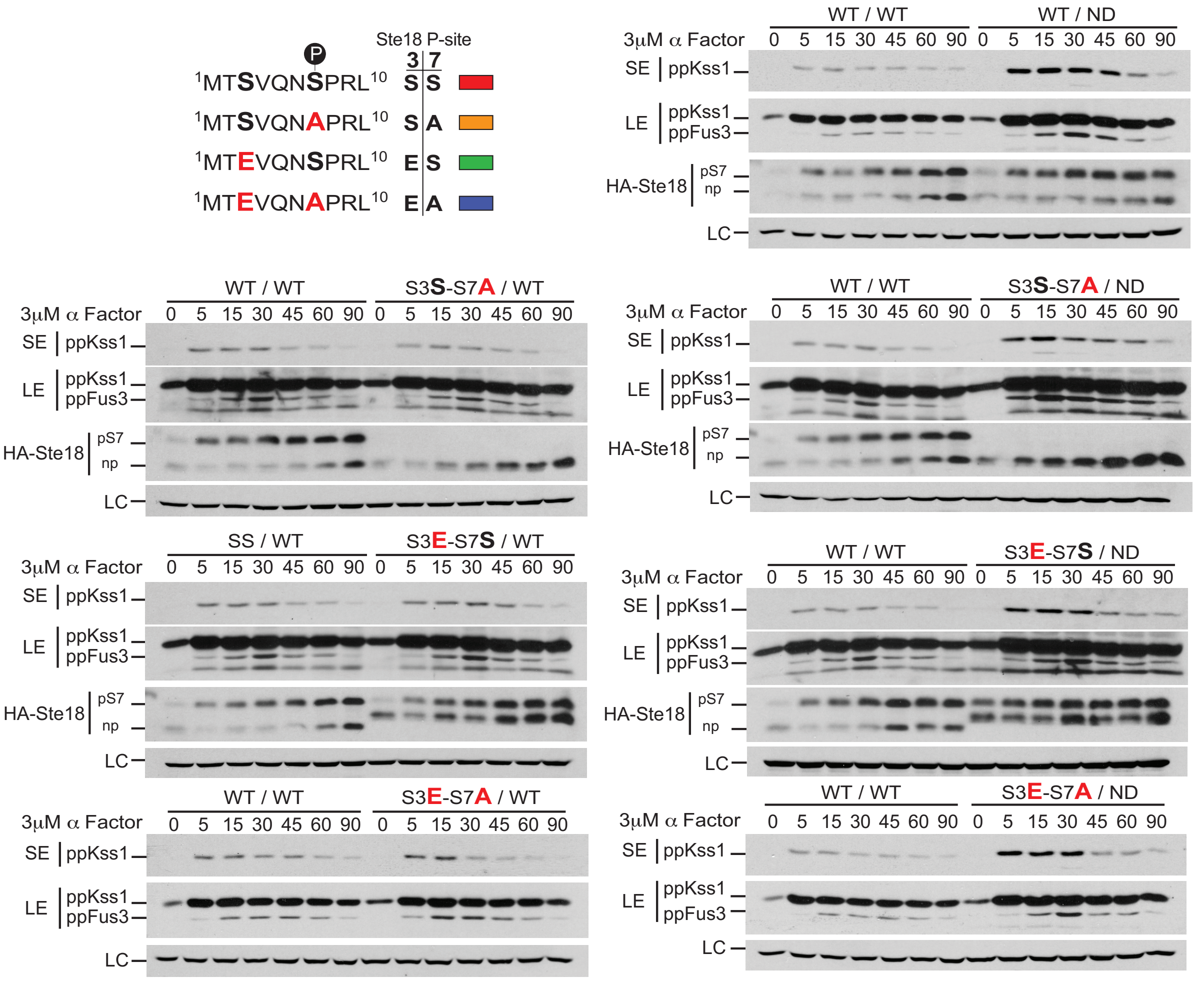


**Figure S4.** Representative immunoblots for activated Kss1 (ppKss1), activated Fus3 (ppFus3), HA-Ste18$^{WT}$ and indicated HA-Ste18 mutants after exposure to 3μM pheromone for the indicated time. SE, short exposure; LE, long exposure; LC, GAPDH loading control.

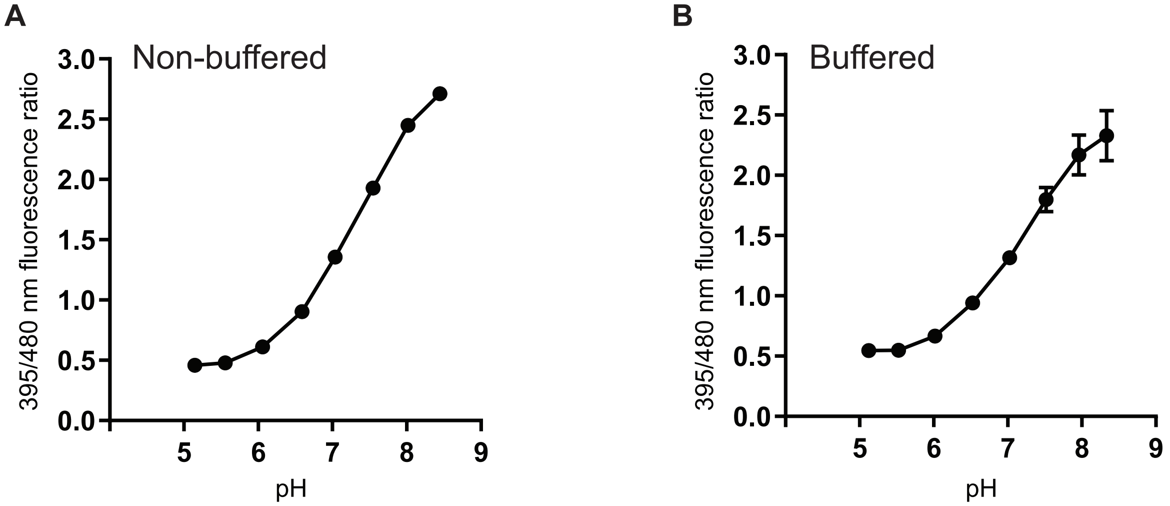


**Figure S5.** pHluorin calibration curves. The ratio of fluorescence intensities emitted upon excitation at pHluorin's two maximum excitation wavelengths (395 nm/480 nm) at different standard pH of cells grown in the absence (A) or presence (B) of buffer. Results are the mean ± SEM (n=4).
